# A longitudinal fNIRS hyperscanning dataset of same generation and intergenerational relationship development and collaboration

**DOI:** 10.64898/2026.06.26.734715

**Authors:** Ryssa Moffat, Emily S. Cross

## Abstract

Social disconnection (i.e., loneliness and social isolation) poses a severe risk to health and is particularly prevalent among older adults. Qualitative and behavioural evidence suggests that intergenerational social interactions can mitigate social disconnection by offering opportunities for adults to form new social connections, yet the neurophysiological and kinematic processes underpinning relationship formation remain poorly understood. This article presents and validates the *InterGenSynchrony Dataset*– a longitudinal, multimodal hyperscanning dataset acquired during a 6-session program, where 30 same generation and 31 intergenerational dyads met as strangers and became acquainted through creative drawing. The dataset includes dyads’ concurrent recordings of brain activity (i.e., fNIRS hyperscanning), motion capture, self-report measures including loneliness, collaborative behavioural scores, drawings (artefacts of collaboration), subjective experiences in text and interview forms. This dataset is optimised for multimodal investigations into the neural, physiological, behavioural, kinematic, and subjective aspects of relationship formation within and between generations in a real-world setting.

## 1 Background & Summary

Social connections are essential for human flourishing ^1,2^. Social disconnection leads to experiences of loneliness and social isolation, which bear serious negative consequences for physical and mental health ^3,4^. As a result, policy makers, researchers and health practitioners are making widespread efforts to help individuals form social connections ^5–7^. These efforts often target older adults, whose risk of loneliness and social isolation is elevated and for whom forming new social connections is more difficult ^8,9^. Indeed, current estimates predict that in 2050, 25% of people living in regions including Europe and North America will be older than 65 years ^10^, and therefore more likely to experience loneliness or social isolation. It is therefore critically pressing, now more than ever, to understanding how adults (young and old) form social connections.

This article describes the *InterGenSynchrony Dataset* ^11^, a longitudinal multimodal dataset optimised for investigating how social connections take shape between adults from the same and different generations. Thirty-one intergenerational dyads (one person 69+ years and the other 18-35 years) and 30 same generation dyads (both 18-35 years) completed a 6-session creative drawing program.

The longitudinal data captured at 6 sessions per dyad (n= 366) in this dataset include: i) simultaneous functional near-infrared spectroscopy (fNIRS) recordings from each dyad member, ii) motion capture coordinates of dyad’s upper body movements during drawing, ii) a solo-drawn drawing per dyad member, two co-drawing drawings per dyad, ii) self-report measures of loneliness, social closeness within dyads, and attitudes towards other generations, iv) behavioural collaborative tasks and utterance counts, v) subjective responses describing the experience in each session. Additional data acquired at a single session per participant includes vi) a self-report measure of empathy, and vii) a ∼20 minute semi-structured interview about the experience of the creative drawing program. To quantify collaborative behaviour based on the co-drawn drawings, we also collected ratings of the collaborative nature of co-drawn drawing from 103 external raters to whom the 6-week program was unknown.

By design, the data acquisition context resembled art-based and intergenerational community programs, both of which offer repeated opportunities for social connection, and are widely supported policy makers, researchers, and health practitioners ^6,12–15^. Evidence from qualitative and behavioural studies shows that art-based activities yield social and physical health benefits for participants of all ages ^16–19^. Notable benefits of repeated intergenerational interactions include improvements in older adults’ physical, social, and cognitive health ^20–22^, as well as reductions in younger adults’ stereotypes towards older adults and aging ^23,24^.

The InterGenSynchrony Dataset fills two critical data gaps. First, it addresses the underrepresentation of older adults in the hyperscanning studies to date. Second, it is the largest longitudinal hyperscanning data set to date. With respect to the underrepresentation of older adults, one study has simulated interbrain synchrony levels of intergenerational dyads involving children, parents, and grandparents using EEG signals ^25^. One other study reports interbrain synchrony in pairs of older adults during competitive tasks ^26^. A recent study examines interbrain synchrony between older adults learning from younger instructors ^27^. Dikker and colleagues’ simulations did not involve real-world hyperscanning data, and Zhang and colleagues’ data are not openly available for reuse. Liang and colleagues’ ^27^ data are openly archived for reuse, but in keeping with the majority of student– teacher hyperscanning studies, only use two younger instructors for all dyads, enhancing experimental control, but limiting the generalizability of the sample. Turning to longitudinal hyperscanning studies, to our knowledge, six hyperscanning studies have recorded INS at multiple sessions and subsequently analysed the aggregated time points, as opposed to reporting changes in INS across sessions ^28–33^. These studies are reviewed in detail in a position piece by Moffat et al. ^34^. Of these studies, only two share data openly for reuse. Ellingsen and colleagues share dyad-level data used to create published figures ^30^, and Francis and colleagues share data from a 3-session study combining neurofeedback and hyperscanning ^33^.

The descriptive analyses presented in this article validate the procedures and illustrate the usefulness of the data for a broader community of researchers in psychology, as well as social and cognitive neuroscientific domains.

## 2 Methods

Section 2.1 details the methods employed in the main longitudinal hyperscanning study. Section 2.2 details the methods employed to collect external ratings of the drawing from the longitudinal hyperscanning study.

### 2.1 Longitudinal intergenerational hyperscanning

#### 2.1.1 Preregistration

##### 2.1.1.1 Preregistered sample size

In our preregistration (https://osf.io/hz6tm), we aimed to collect useable data from a minimum of 30 same generation and 30 intergenerational dyads between November 2023 and June 2024. We selected this minimum to maximise our financial, temporal, and human resources while obtaining a data set optimised to address numerous research questions ^35^. As preregistered, data collection began on November 15, 2023 and ended on June 18, 2024 (Figure 1A).

**Figure 1.**
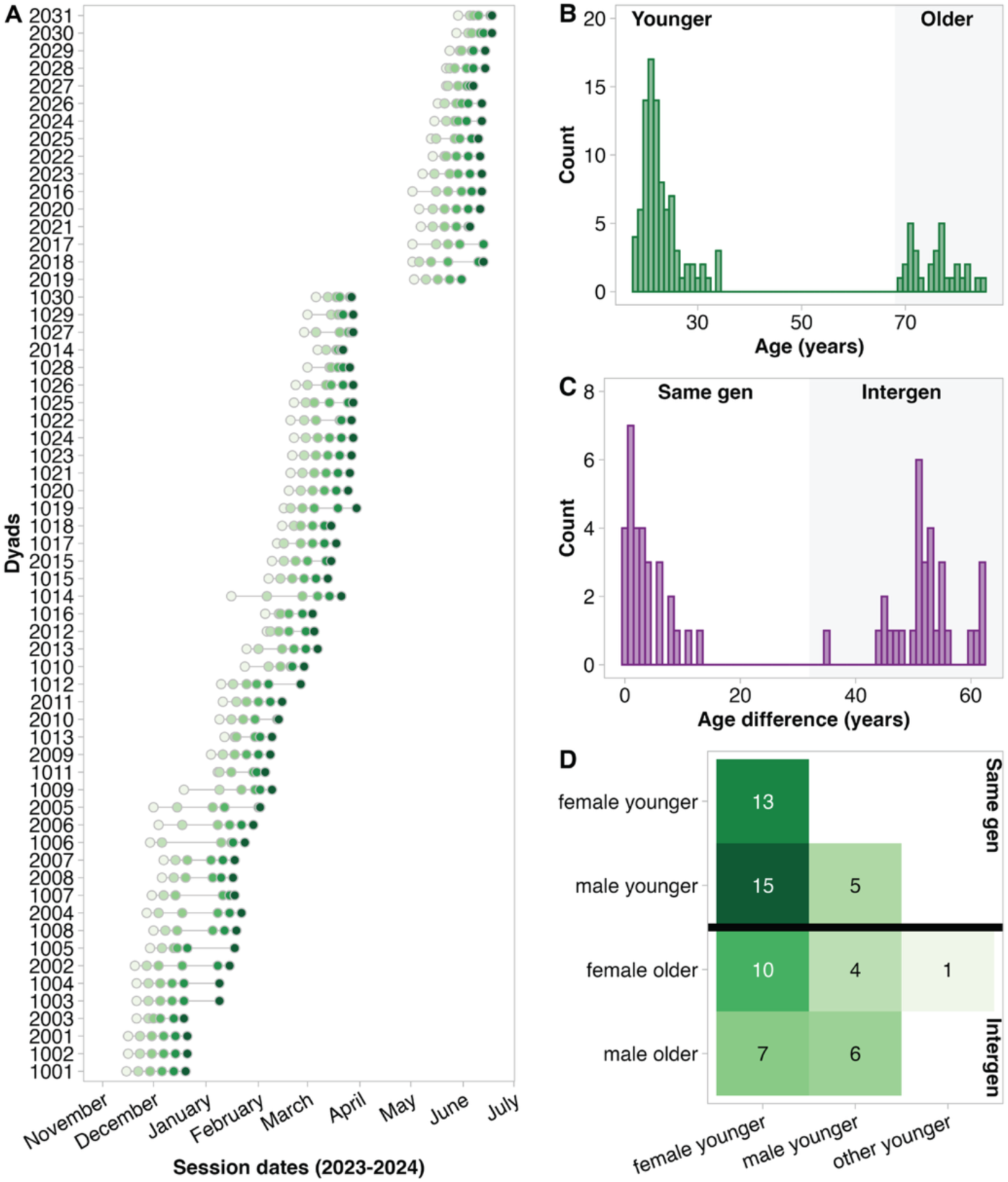
A) Schedule of sessions. B) A histogram of participants’ ages. C) A histogram of age difference within dyads. D)The number of dyads for each of the eight dyad pairings in this sample.

##### 2.1.1.2 Deviations from preregistration

We preregistered that older adults would be 70+ years old. However, due to the challenge of recruiting older adults, we were concerned about achieving the preregistered minimum of 30 intergenerational dyads. We therefore included one 69-year-old participant.

#### 2.1.2 Participants

We recruited 122 community-dwelling participants from Zurich, Switzerland. Of these, 31 were older adults (aged 69–85 years) and 91 were younger adults (aged 18–34 years; Figure 1B). We assigned participants to intergenerational dyads (n = 31) and same generation dyads (n = 30) based on availability for sessions (e.g., matching people available at the same day/time for 6 consecutive weeks). Age differences within dyads are visualised in Figure 1C. Due to the logistical challenge posed by recruiting for and scheduling a multi-session study of this scale, gender pairings were not prioritised (see 8 possible gender pairings in Figure 1D). Intergenerational dyads consisted of an older adult (range: 69–85 years, mean age: 76 ± 4 years; 18 female, 13 male) with a younger adult (range: 18–34 years; mean age: 24 ± 4 years; 19 female, 11 male, 1 other). Same generation dyads consisted of two younger adults (range: 18-34 years; mean age: 22.5 ± 3.6 years; 36 female, 24 male). All dyads started the study as strangers. To be included, participants had no known history of neurological or psychological disorders (e.g., stroke, concussion, ADHD, autism, schizophrenia, depression) and were fluent in German. No further demographic information was collected. Seven dyads (3 same generation, 4 intergenerational) attended the initial session, then unenrolled. In accordance with our preregistration, their data are not included in the analysis.

Younger adults were primarily recruited through ETH Zurich’s DESCIL student pool. Older adults were primarily recruited via the University of Zurich’s Healthy Longevity Centre, clubs and organizations for senior citizens (e.g., theatre groups and choirs). We also recruited using flyers and social media.

#### 2.1.3 Ethics statement

Ethical approval was obtained from the Ethics Committee of the Canton Zurich (Ref: 2023-01073). The study was conducted according to the Declaration of Helsinki. We obtained written informed consent from all participants.

#### 2.1.4 Procedure

Dyads attended six sessions, with approximately one week between sessions (Figure 1A). All sessions had the same structure (Figure 2A). Sessions were led in High German (Hochdeutsch, as opposed to Schweizerdeutsch) by a female lead researcher who facilitated sessions following a script (available on OSF: https://osf.io/zms2v) and answered participants’ technical and scientific queries. A second female researcher was present to aid with technical steps and did not speak unless directly addressed. Both researchers politely deflected participants’ social inquiries. To ensure that all participants had equal amounts of contact with the experimenters, a research assistant was responsible for scheduling sessions and sending participants reminders about upcoming sessions. All sessions were held in a ∼16 m^2^ room, arranged as shown in Figure 2B-C.

**Figure 2.**
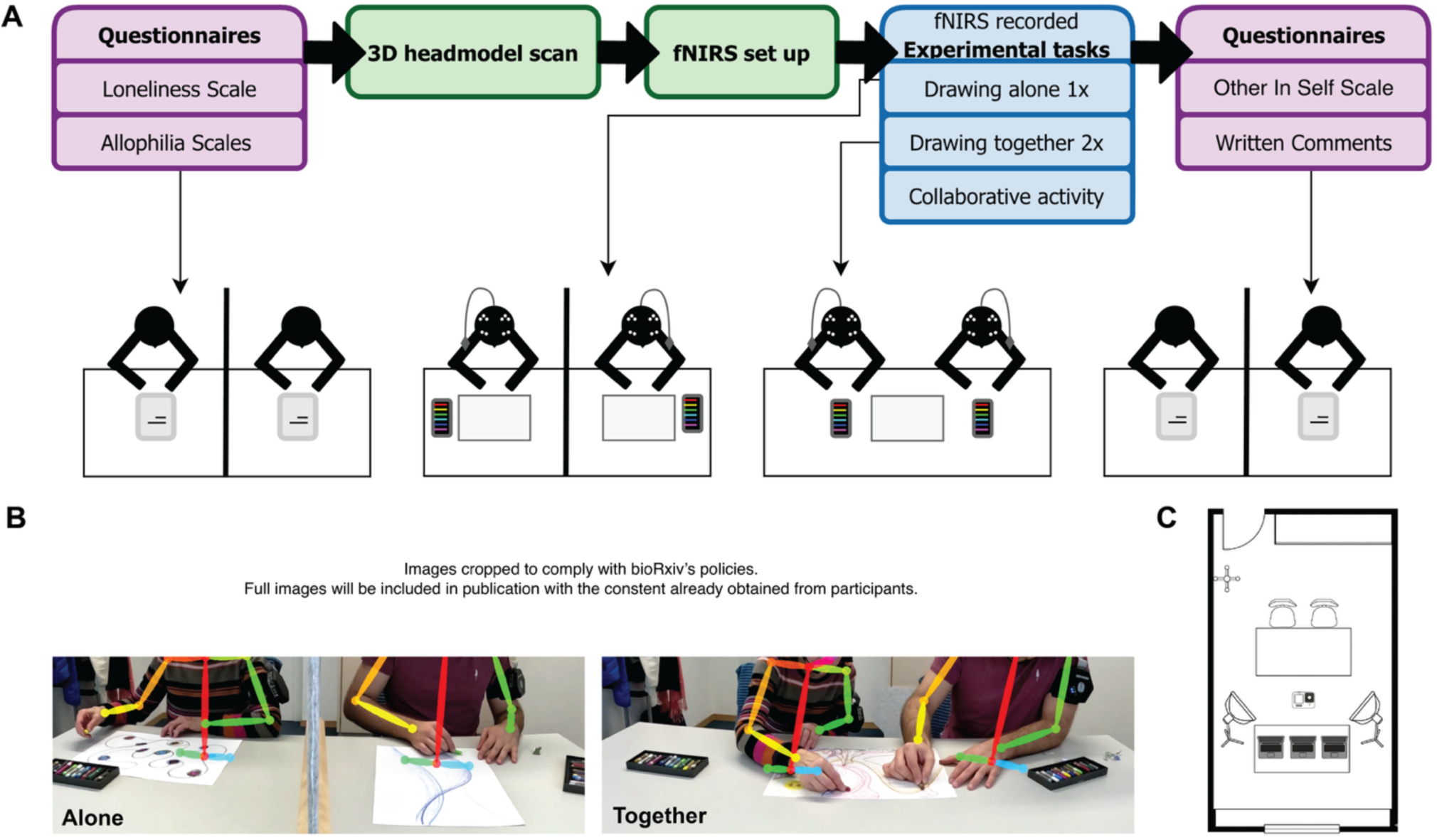
A) A schematic showing how each session was structured. At the first session, participants also completed an empathy questionnaire at the beginning of the session. At the end of the final session, participants were interviewed by the lead researcher about their experience in the study. B) Still photographs from video recording showing an intergenerational dyad drawing alone (left) and together (right). Photographs captured by Ryssa Moffat. Both participants depicted in these photographs have provided written consent for their likeness to be published in open-access online research outputs, such as this dataset description. C) Floor plan of room, in which sessions were held. Floor plan image adapted using Gemini.

### Self-report measures and technical setup

At the beginning of each session, dyads completed self-report measures of loneliness and attitudes towards generations (allophilia) on an Apple iPad Pro 10, with a felt divider (50 cm high) between participants providing privacy. Self-report measures were administered via RedCap. When both participants finished, the researchers placed the fNIRS caps on participants’ heads. Next, the second researcher recorded 3D models of each participant’s head with the cap on (no optodes attached) using a Sensor Structure Pro (model SA39) attached to an Apple iPad Pro 10. The lead researcher instructed each participant to sit very still while the second researcher circled the participant with a 3D scanner, recording the 3D head model. Next, the lead researcher reminded participants that they could converse at any time except while drawing. Then, the researchers attached optodes to the participants’ fNIRS caps, ensuring that optodes came in direct contact with the scalp by parting hair. The researchers turned the fNIRS devices on, paired them with the recording laptops (n = 2), and verified the fNIRS signals for best possible calibration. The researchers initiated the video recording, which was captured by a GoPro11 Black camera positioned on a tripod, to obtain a direct view of participants (Figure 2B-C).

### Drawing and collaborative activities (during fNIRS recording)

Next, participants drew three drawings with oil pastels, without talking. The allotted time per drawing was 5 minutes. The lead researcher instructed the participants that they would draw three times, each time receiving 5 minutes to draw anything they liked, and that they would complete the first drawing alone and the second and third drawings together with their partner on a single piece of paper (A3, 170gsm). In preparation for the first drawing (alone), a researcher laid a separate piece of paper in front of each participant, and a felt divider was positioned between participants, ensuring that they could not see each other. The lead researcher specified that participants should not talk during the 5 min of drawing time, provided no further instructions (e.g., no suggestions for motifs). The lead researcher then simultaneously said “start” and pressed a keyboard button on the experimental laptop that sent a trigger to the recording computers and started a 5-minute timer. The timer was visible on the laptop screen. At the end of 5 minutes, the lead researcher said “stop”, the researchers retrieved the drawings and removed the felt divider. A researcher then laid a single piece of paper on the table. The lead researcher instructed participants that they would have 5 minutes to draw together and should not talk during this time, paused then said “start” and pressed the trigger/timer button. This was repeated for the second instance of drawing together. The lead researcher systematically alternated between two German words for together to ensure that participants self-selected their level of collaboration (“zusammen” and “gemeinsam”, which translate to “together” and “with one another” respectively). In cases where participants conversed to plan their drawings, the lead researchers waited for a natural pause to begin the 5-minute timer. If a dyad had not conversed during sessions 1-3, at the fourth session, the lead researcher gave a prompt to signal that conversation is allowed before beginning the timer for the first instance of drawing together. The prompt was “abgesprochen?”, which translates to “agreed?” or “all decided?” and implies prior conversation.

Following the three drawings, the participants completed a short collaborative activity. Five different tasks were selected (described in detail in Section 2.1.4.6), with one activity repeated at the first and last session to measure changes in collaboration, and the remainder varied to maintain participants’ engagement in the study. The lead researcher also said “start” and pressed the trigger/timer button to initiate the collaborative task.

### Self-report measures, subjective experiences, and technical dismantling

After the collaborative activity, the researchers stopped the fNIRS recordings and placed the felt divider between participants to provide privacy. Still wearing the fNIRS caps, participants completed a self-report measure of social closeness (OIS) and provided a written subjective description of their experience in the session. Both the self-report measures and the subjective experience question were administered via RedCap using Apple iPad Pro 10s. When the first participant finished, a researcher retrieved the iPad, disconnected the optodes from that participant’s fNIRS cap, and then removed the fNIRS cap. When the second participant finished, the other researcher removed the felt divider and subsequently removed the fNIRS equipment from the second participant’s head. The lead research thanked the participants for their attendance and the participants departed.

### Interview after final session

After the final session, the lead researcher held a semi-structured interview with each participant individually to gain qualitative insights into the participant’s experience.

#### 2.1.5 Measures

##### 2.1.5.1 Self-report measurements

###### Empathy

At the beginning of the first session, participants completed the 16-item *Saarbrücker Persönlichkeitsfragebogen zur Messung von Empathie* ^36^, which is a German version of the *Interpersonal Reactivity Index* ^37^.

###### Loneliness

Participants completed the *6-Item Scale for Overall, Emotional, and Social Loneliness* ^38^ at the beginning of each session.

###### Attitudes toward other generations

The *Allophilia Scale* ^39^ is typically used to characterise participants’ attitudes toward an outgroup (e.g., people of another ethnicity or political leaning). We adapted this 17-item scale in three variants: a) “younger people”, “older people”, and “people my age”. At the beginning of each session, all participants completed the scale with “people my age”.

Next, older adults completed the scale with “younger people” and younger adults completed the scale with “older people”. By having participants record responses for “people my age” and the age group they did not belong to, a baseline correction of attitudes towards other generations is possible.

###### Social closeness

Participants indicated how socially close they felt to the other member of their dyad using the single-item *Inclusion of Other in the Self Scale* ^40^ at the end of each session. This measure uses Venn diagrams with incremental increases from no overlap to complete overlap as visual analogue representation of social relationships.

##### 2.1.5.2 fNIRS recordings

We recorded fNIRS signals using two Cortivision Photons Cap (Cortivision sp. z o.o., Lublin, Poland) and Cortivew software. Each fNIRS device transmitted the recorded signal to a separate recording laptop (n = 2; right and left laptop in Figure 2C). A third laptop (middle laptop in Figure 2C) was used to synchronise the fNIRS recording by sending triggers from PsychoPy ^v. 2022.2.5;,41^ that marked the beginning of each drawing and the collaborative activity. The laptops were cabled together with a powered switch (TP-Link 5-ports) and ethernet cables to minimise the risk that a trigger might not be received. Trigger 1 corresponds to drawing alone, trigger 2 to the first instance of drawing together, trigger 3 to the second instance of drawing together, and trigger 3 to the collaborative activity.

Each fNIRS device had 26 optodes. This included 10 detectors and 16 LED sources emitting wavelengths of 760 and 850 nm. Four of these sources are designed to be used in short channels for measuring from superficial tissue. The sample rate was between 4 and 6 Hz. To position the optodes over bilateral IFG and TPJ, we searched the terms ‘ifg’ and ‘tpj’ in the Neurosynth database (https://neurosynth.org/). We located the coordinates of the brain area in the search results with the highest activation in each the left and right hemisphere to use as the ‘centre’ of our ROIs. These were the following: *left IFG:*-48, 22, 8; *right IFG:* 50, 22, 16; *left TPJ:*-56,-54, 20; and *right TPJ:* 56, - 52, 18. We then positioned optodes in the 10-5 EEG positions ^42^ around these regions, guided first by the fOLD toolbox ^43^ with validation using optode digitization and 3D head models (Figure 3).

**Figure 4.**
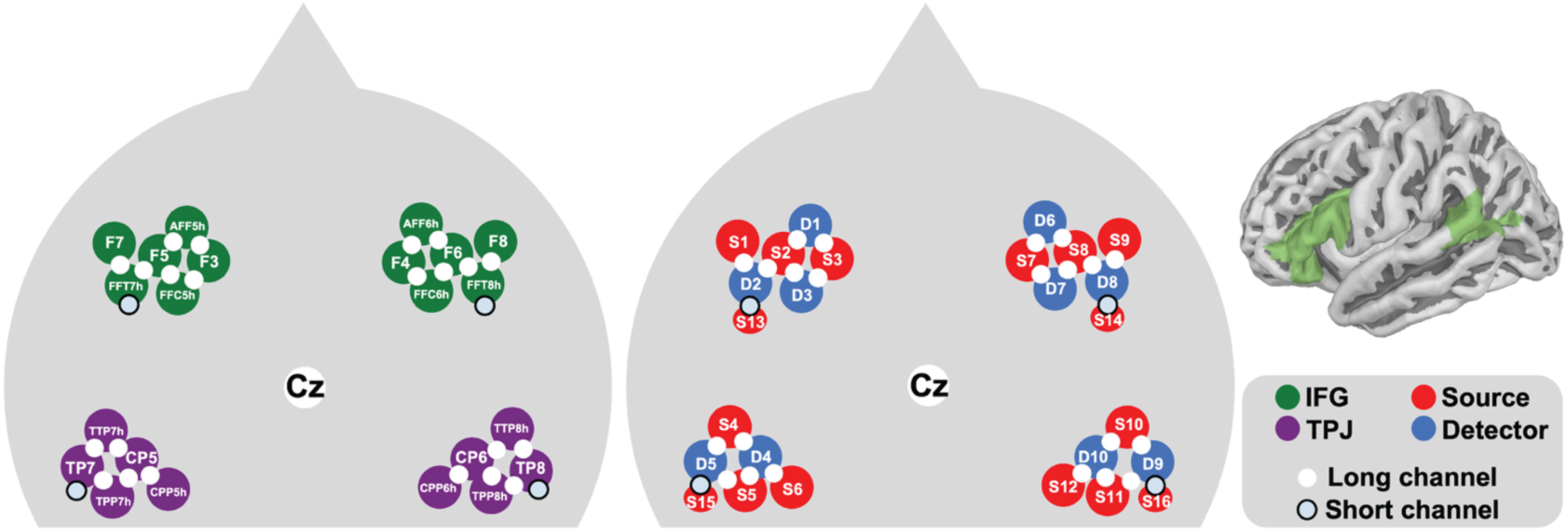
Arrangement of optodes. The left head shape shows the 5-10 positions of each optode within the four ROIs (bilateral IFG and TPJ), as well as short and long channels. The right head shape shows the source and detector numbers that make up each short and long channel. With Cortivision Photon Cap fNIRS device, short channels are created using specific sources (S13-S16), as opposed to more typical specific detectors. On the inset brain, the green shading indicated the IFG and TPJ cortical regions.

###### Digitisation of optodes

The researchers positioned the fNIRS cap on participants heads according to fiducial markers. Before attaching optodes, the researchers recorded a 3D image of each participant’s head using Structure Sensor Pro (model SA39) attached to an Apple iPad Pro 10. We processed the 3D head scans using MATLAB (v. 2023b) and the Fieldtrip toolbox ^44^. First, we assigned labels for each detector, sensors, and fiducial reference point to the 3D head scan and then co-registered these to MNI space to obtain coordinates. With these coordinates, we computed the inter-optode distance for each channel, the coordinates of the point of measurement between pairs of optodes that formed channels, and distance from the point of measurement to the centre of the ROI.

##### 2.1.5.3 2D motion capture

We video-recorded participants’ upper body movement while they drew alone and together, as well as during the collaborative activities. Videos were recorded at a rate of 60 frames per second with a GoPro11 Black positioned ∼1.25 m in front of participants on a tripod, with the camera raised to ∼1.25 m from the ground. We estimated both dyad members’ poses in each frame using OpenPose ^45^. Specifically, we estimated the x and y coordinates in the video frame of each dyad member’s nose, neck, as well as their right and left eyes, ears, shoulders, elbows, and wrists.

We estimated both dyad members’ poses in each frame using OpenPose ^45^. Specifically, we estimated the x and y coordinates in the video frame of each dyad member’s nose, neck, as well as their right and left eyes, ears, shoulders, elbows, and wrists. OpenPose returns coordinates with confidence levels for each identified joint ^45^. To ensure that all coordinates were assigned to the correct dyad member, we used in-house code (https://osf.io/akmqw/files/4rn5f) to match each dyad member’s neck (and by extension, further joints) across frames. For frames where the neck was identified with a confidence below 0.5, we set all coordinates to *NA* (not available).

##### 2.1.5.4 Drawings

Participants drew on A3 170gsm paper. Each participant had access to a separate set of 12 oil pastels (Caran D’Ache Neopastel). After the session, the researchers labelled each drawing with the participant or dyad ID and session using a ball-point pen and then photographed the drawings using an Apple iPhone Pro 12. Scanning was not possible because the oil pastel would transfer from the paper to the scanner. A research assistant straightened the photos of drawings, adjusted the white balance, and removed the dyad ID from the drawing using Adobe Photoshop. The JPG files were converted to TIFF to ensure lossless archiving.

##### 2.1.5.5 Experimenter ratings of simultaneous drawings

To quantify simultaneous drawing and turn-taking behaviour while dyads drew together, the lead experimenter watched a timer counting down from 5 minutes and estimated the percentage of the 5 minutes that dyads drew simultaneously in increments of 10%. The percentage was then divided by 10, yielding a score between 0 and 10. Zero indicates that the dyad engaged exclusively turn-taking (i.e., 100% turn-taking and 0% simultaneous drawing). Ten indicates that the dyad engaged only in continuous simultaneous drawing (0% turn-taking and 100% simultaneous drawing).

##### 2.1.5.6 Collaborative behaviour

A research assistant, fluent in Swiss German, reviewed the video recordings to behaviour and score the collaborative tasks. The research assistant then recorded the number of utterances produced by dyads during collaborative tasks where talking was permitted but not directly scored (i.e., sessions 1, 3, and 6). The research assistant also recorded whether or not dyads engaged in verbal planning before drawing together, and if a dyad engaged in planning, the number of utterances produced.

###### Session 1 and 6

The lead experimenter placed both participants’ oil pastels (12 per participant, 24 total) in a pile in the center of the table and then instructed participants to sort the pastels into their respective boxes. The time (in seconds) to the final crayon being played in its box, and the accuracy of the sorted crayons were recorded.

###### Session 2

Participants completed a dyadic verbal fluency task. They took turns saying nouns (proper nouns not permitted) that start with the letter “p” for 1 minute. The total number of High German and Swiss German common nouns was counted.

###### Session 3

The lead researcher placed the pieces of a 54-piece jigsaw puzzle (Djeco Giant Puzzle, image of animals in a forest; piece dimensions: ∼ 8 x 8 cm) in the centre of the table and instructed the participants that they had 6 minutes to assemble as many pieces as possible and that talking was permitted. The number of assembled puzzle pieces was counted. In the case that participants assembled all pieces in less than 6 min, the remaining number of seconds from 6 min was divided by 6.66 (the average number of seconds per piece if completed in 6 min; 360 s/54 pieces = 6.66 s) and added to a score of 54.

###### Session 4

Participants completed a dyadic divergent thinking task. The lead researcher instructed participants to name as many uses for a lumpy blue rubber ball (⌀ 5 cm Gaiam foot massage ball) as possible in 2 minutes and that turn-taking was not required. The number of unique uses was scored.

###### Session 5

Participants completed a dyadic verbal fluency task. They took turns saying verbs that starting with the letter “p” for 1 minute, with the aim of saying as many as possible. The total number of High German and Swiss German common verbs was counted.

##### 2.1.5.7 Subjective experiences

###### Written responses at each session

At the end of each session, participants had the opportunity to describe their experience subjectively, either by filling in a text box on an iPad or writing by hand on paper. Participants received the following open-ended prompt: “We would like to better understand your experience today. Please describe how you perceived the art activities today”. Research assistants redacted personal information and emojis from the responses.

###### Semi-structured interview after final session

After completing the final session, the lead experimenter conducted a semi-structured interview with all participants (prompts in Table 1). She asked participants to speak High German during the recording (though some participants naturally reverted to Swiss German). Interviews were recorded in.m4a format using an Apple iPhone Pro 12 and the Voice Memos app, then transcribed using Trint (https://trint.com/). Two research assistants, both fluent in Swiss German, reviewed transcripts alongside the audio recordings to ensure correctness and redacted personal information.

**Table 1.** Using these questions, the lead researcher conducted a semi-structured interview with each participant individually at the end of the final session.

| Questions |  |
| --- | --- |
| 1. | Please describe your partner in 3 words. |
| 2. | Do you have any general comments about the experiment? |
| 3. | Did anything change for you, from the first to the last session? For example, while drawing or in the interaction? |
| 4. | During the experiment, what did you find to be a) the best, b) the worst, c) the most surprising? |
| 5. | How did you perceive your contribution to the co-drawing? And during the other collaborative tasks? |
| 6. | Did you feel valued by your partner? What signs did you notice? |
| 7. | Did you learn anything about yourself during the experiment? |
| 8. | How did you choose the themes/motifs while drawing together? |
| 9. | Were the themes more often realistic/figurative or abstract, like modern art? Why was this? |
| 10. | Did you ever add details to your partners' figures? Why/why not? What would have happened if you had of? |
| 11. | How did you perceive the role of the experimenters leading the study? How did we influence your experience? |
| 12. | Were you in contact with your partner outside of the study? How long in minutes on average? |
| 13. | Do you think your experience would have been the same if you were assigned a partner from another generation/from your own generation? |
| 14. | At the end of each session, you answered a question with many overlapping circles. Did you have any thoughts about that question? |
|  | Other comments? |

### 2.2 External ratings of drawings

#### 2.2.1 Participants

To obtain ratings of drawings produced by the main sample (described further in 2.3.1), we divided the 726 drawings into 6 sets of 121 drawings. We aimed to collect 20 ratings per drawing by having 20 people rate each set of drawings. In total, we recruited 148 new participants via Prolific after the main study concluded. To be included, participants had to be over 18 years old, proficient English speakers, residents of the United Kingdom, and not yet have completed our rating experiment (i.e., participants could only rate one set of drawings). Beyond these criteria, we did not collect demographic information from these participants. To ensure response quality, we only accepted participants who had previously completed a minimum of 50 tasks on Prolific with a 100% approval rate. Of the 148 recruited participants, 45 were excluded for failing to respond to all attention checks correctly (i.e., by moving all sliders all the way to the right of the sliding scale when prompted). Our analyses thus include 103 raters, where each drawing was rated by between 14 and 20 individuals (mean = 17.23, SD = 1.49).

#### 2.2.2 Ethics statement

Ethical approval was obtained from the Ethics Committee of ETH Zurich (Ref: 2024-N-265). The study was conducted according to the Declaration of Helsinki. All participants provided written informed consent.

### 2.3 Procedure

#### 2.3.1 Ratings of drawings

Ratings were collected using a Qualtrics survey, in which drawings and ratings were presented as shown in Figure 5. Before beginning the survey, raters were informed that they would view drawings that had been created by two people at once on a single piece of paper, and that they would rate the drawings on three scales. After providing written consent, raters viewed and rated the drawing in Figure 5 to become familiar with the experiment format. Then, each rater viewed 121 drawings in a randomised order, one at a time. For each drawing, raters rated the degree of harmony between the elements of the drawings (on a sliding scale from 0 “entirely independent” to 100 “entirely coordinated”) and the use of space (on a sliding scale from 0 “divided” to 100 “unified”).

**Figure 5.**
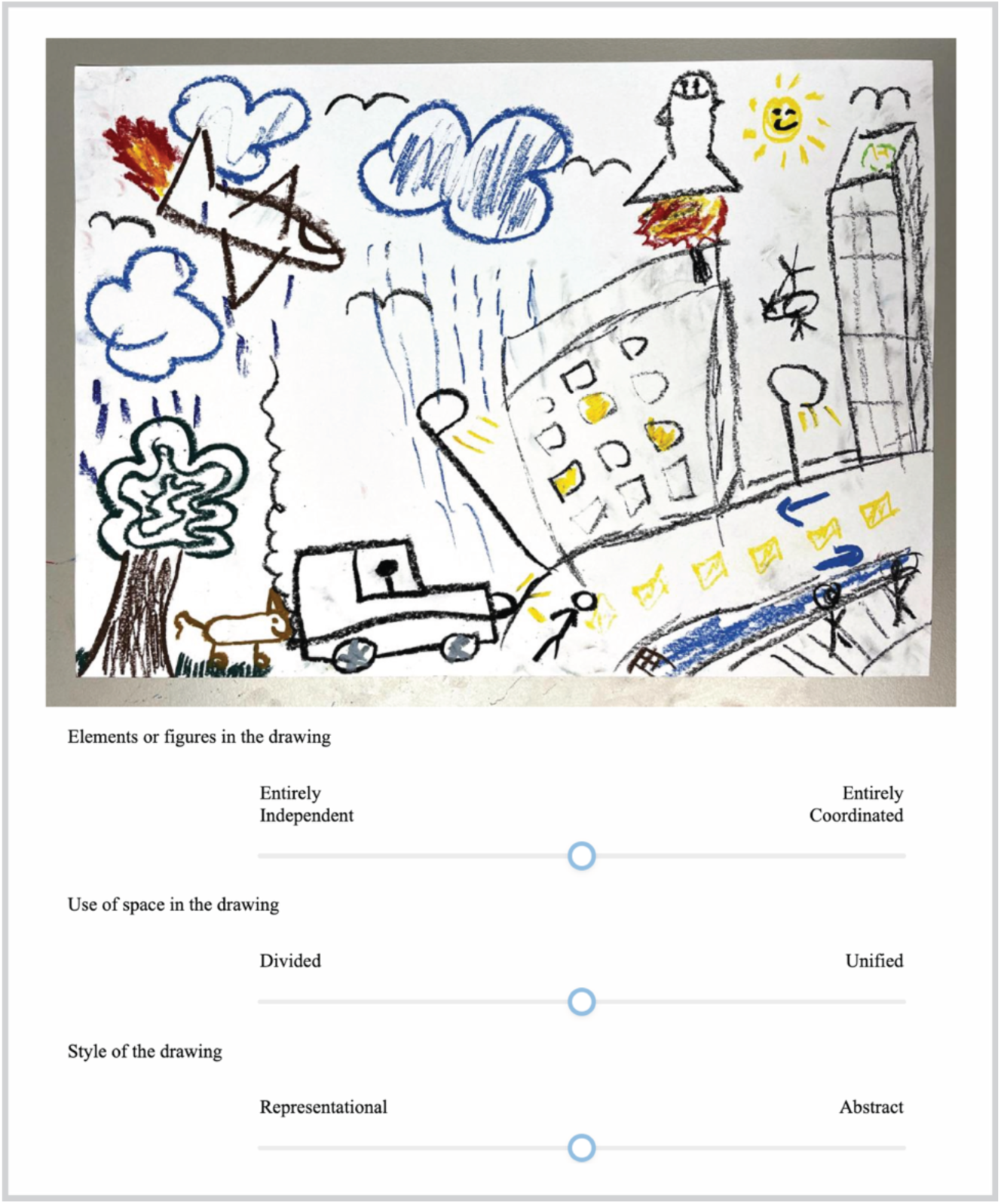
The presentation of drawings and rating scales using Qualtrics.

## 3 Data records

The InterGenSynchrony Dataset ^11^ was released on July 10th, 2026 on OpenNeuro (https://openneuro.org/datasets/ds008192). The InterGenSynchrony Dataset is shared under a CC0 licence. The dataset is formatted according to BIDS (Brain Imaging Data Structure) guidelines and adapted for longitudinal dyadic data as recommended by a BIDS maintainer ^46^.

The structure of the InterGenSynchrony Dataset is visualised in Figure 6. As per BIDS structure, the fNIRS brain imaging data are in the main directory. All other data modalities are in the “sourcedata” directory. Inside “sourcedata” there are directories for the non-brain imaging data from the main study (“hyperscanning”) and the external ratings of drawings (“ratings”).

**Figure 6.**
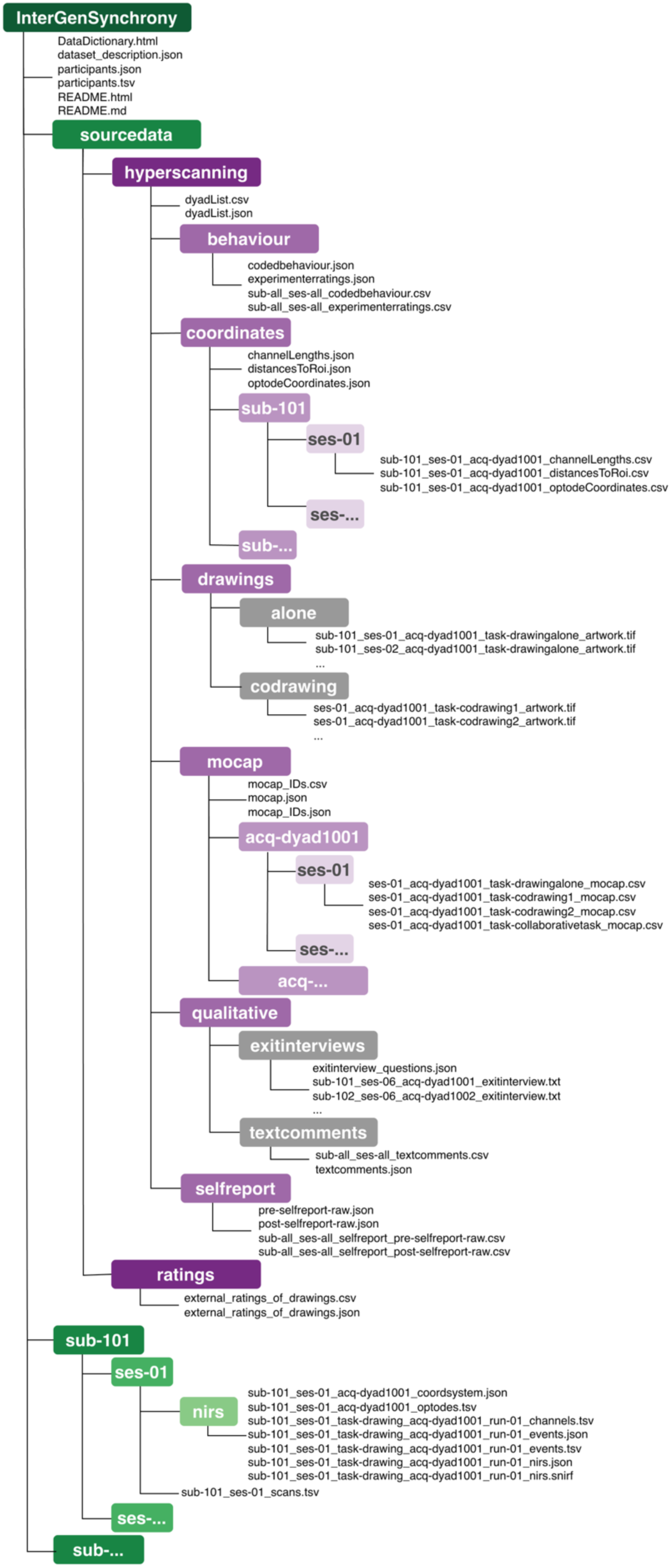
The BIDS-compatible structure of the dataset. To describe the content of CSV files, we provide an easy-to-read Data Dictionary, as well computer-readable JSON files, which are named to match the relevant CSV files.

## 4 Technical Validation

For the fNIRS and 2D motion capture data, we report measures of signal quality. For the self-report data, we assess the internal consistency of the multi-item measures. For the self-report, behavioural, and rating data, we visualise the distributions to characterise the available data. No validation procedures were conducted with the qualitative data. Finally, where relevant, we refer readers to existing analyses of each data type.

### fNIRS

We computed the scalp-coupling index for frequencies between 0.7 and 1.35 Hz, per channel. As illustrated in Figure 7A, 88% of channels have an SCI ≥0.5 and <1. We recommend excluding channels with SCIs <0 (or preferably a higher threshold) and >1 from analyses, as these signals were recorded with defective sensors, which were quickly identified and replaced as rapidly as possible.

**Figure 7.**
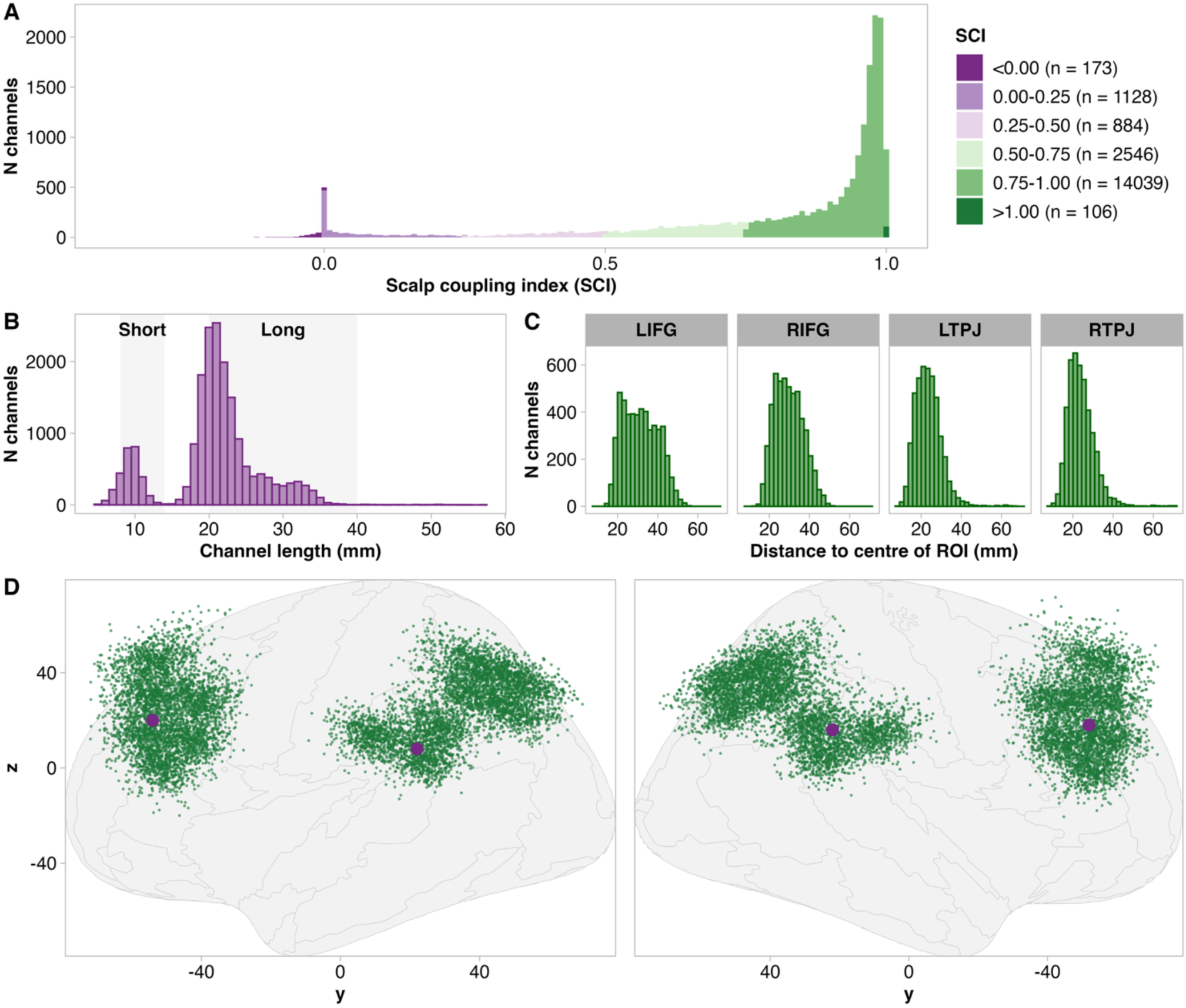
A) Histogram showing distribution of scalp-coupling indices (SCIs) of all fNIRS channels. Colours correspond to 0.25 SCI bins. Channels with SCIs <0 and equal to 1 are included in separate bins, as these were the result of defective optodes, which were quickly identified and replaced. B) Histogram showing the distribution of channel lengths in mm. In this figure, channels with source–detector distances of 8-14 mm are labelled as short channels and channels with source– detector distances of 20-40 mm are labelled as long channels. C) Histograms per ROI showing the distance of the estimated point of measurement of each channel from the centre of the ROI in mm. D) A 2D representation of the digitised location of all recorded channels (green dots) and the centres of each ROI (purple) in MNI coordinates.

Additionally, all channels were visually inspected for cardiac oscillations by a research assistant. Channels without cardiac oscillations or with major recurrent motion artifacts are flagged as “bad” in the recording-specific files detailing channel information (i.e., channel.tsv files).

In addition to SCI values, Figure 7 also provides information on channel lengths, distances of channels from the centre of the respective ROIs, and the coverage of the cortex. We note that the distances from the centre of the ROIs are variable and, in many cases, large. We recommend that specific position of each channel be considered individually, rather than the assigned ROI. This could involve selecting the closest channel of adequate quality, setting a threshold for an acceptable distance from the ROI centre and selecting multiple channels, or identifying the anatomical locations that correspond to channel-position before conducting statistical analyses ^47,48^.

Further confirming the validity and usability of this dataset, we conducted an empirical analysis of this data ^47^. One component of this analysis was a comparison of interpersonal neural synchrony levels during independent and collaborative drawing. Previous studies on collaborative and cooperative interactions report heightened interpersonal neural synchrony during collaborative, relative to independent behaviour ^49–55^. Consistent with these findings, we found that INS was greater during collaborative drawing than during independent drawing in four pairs of brain regions, implicating both left and right IFG and TPJ ^56^. Based on this demonstration, these fNIRS data are aligned with the theoretically expected outcomes.

### 2D motion capture

OpenPose returns coordinates with confidence levels for each identified keypoint ^45^. Figure 8 shows the measure of central tendency (mean and standard deviation) for confidence levels per keypoint, as well as the proportion of cases where OpenPose was unable to recognise a keypoint. Both metrics are calculated based on a maximum of 11906819 recorded observations per keypoint. Except for the left and right ears, all key points had a mean confidence level > 0.7. Similarly, except for the left and right ears, the proportion of unrecognised keypoints is minimal. These values align with those in other high-quality 2D motion capture datasets ^57^. The videos show that the left and right ears were typically occluded by head turns. Caution should be taken when anlysing the time series of the left and right ears.

**Figure 8.**
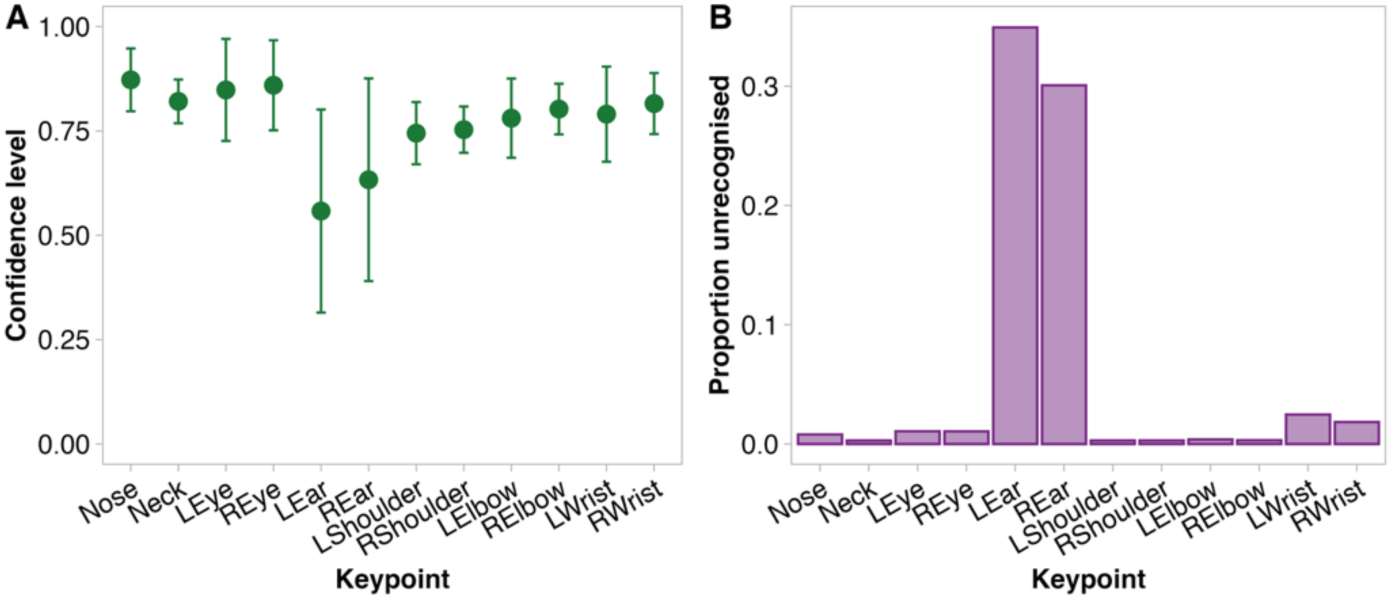
A) Mean and standard deviation of confidence levels per OpenPose keypoint. Confidence values range from 0 to 1, where 1 indicates maximum confidence, and 0 indicates the lowest confidence that the coordinates returned for a specific keypoint are accurate. B) The proportion of frames in which each keypoint was not recognised or where the confidence level for the neck was <.5.

### Self-report

Distributions of self-reported measures are visualised in Figure 9. We assessed the internal consistency scales with multiple items using Cronbach’s alpha ^58^. The 16-item empathy scale showed good reliability (*α* = 0.75, 95% CI [0.72, 0.77]). The 6-item loneliness scale showed acceptable reliability (α =.67, 95% CI = [0.62, 0.69]). Each of three allophilia 17-item scale variants showed excellent reliability (*α* _same-aged people_ = 0.94, 95% [0.93, 0.94], *α* _older people_ = 0.94, 95% CI = [0.92, 0.94], *α* _younger people_ = 0.93, 95% CI = [0.92, 0.94]). With the exception of empathy, these scores have been reported in an empirical analyses ^59,60^.

**Figure 9.**
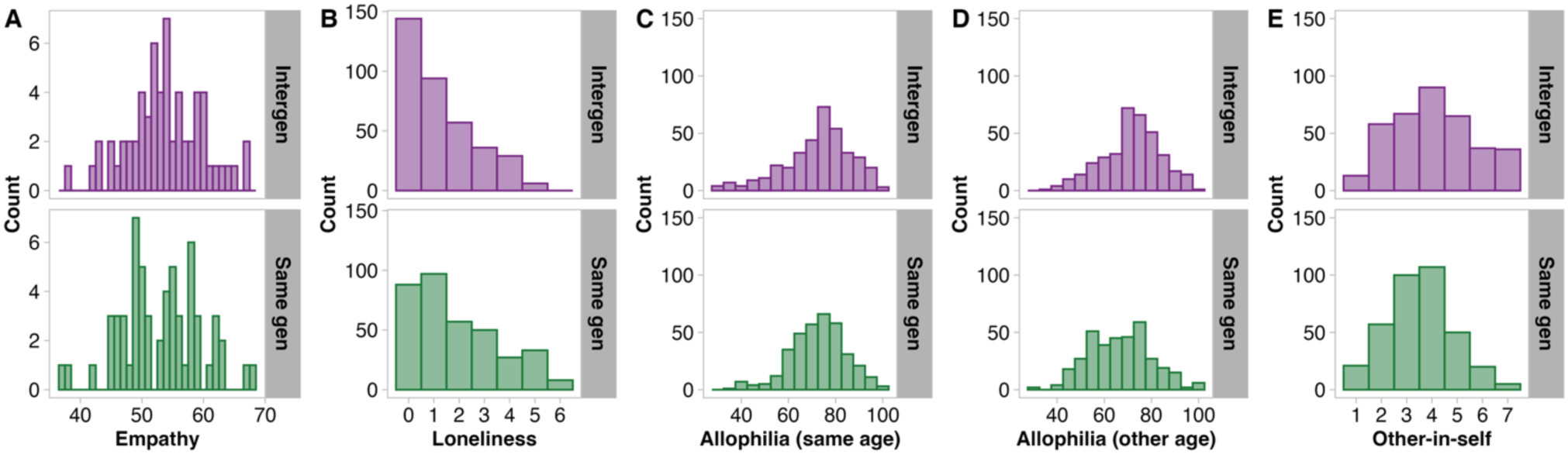
Histograms per group showing distributions of A) self-reported empathy, B) loneliness, attitudes toward C) one’s own and D) the other age group (allophilia), and E) social closeness (other-in-self). Empathy scores were only recorded before the first session. All other measures were recorded at every session. Intergen = intergenerational group, Same gen = same generation group.

### Collaborative behaviour

The distributions of scores for each collaborative task are presented in Figure 10A. The distributions of collaborative behaviour, such as simultaneous drawing and verbal planning immediately before and while drawing together, are shown in Figure 10B-D. The puzzle task scores ^59^ and experimenter ratings of simultaneous drawing ^60^ have been reported elsewhere.

**Figure 10.**
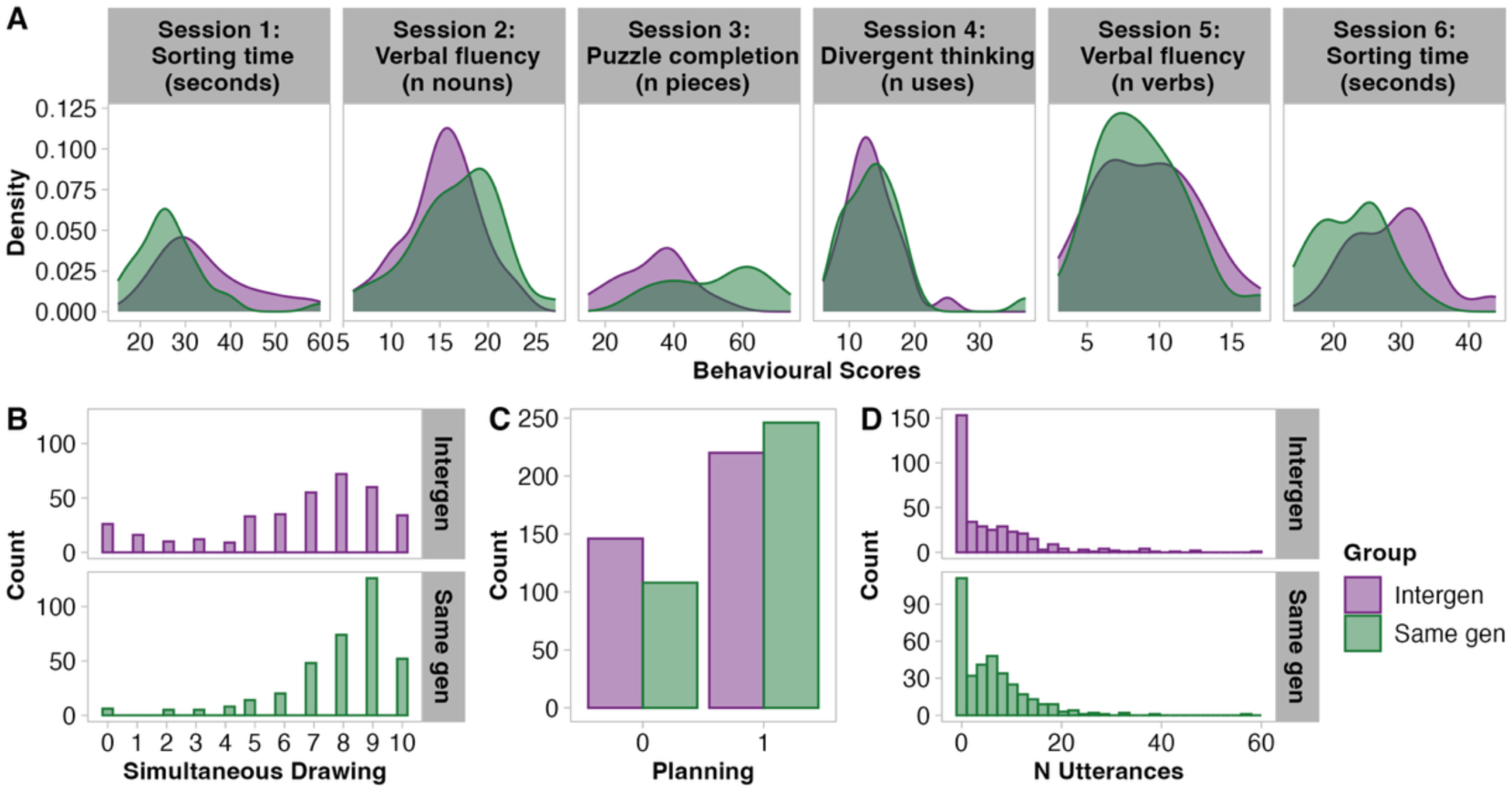
A) Distributions of scored behaviours recorded during collaborative activities. B) Histograms of simultaneous drawing, as rated by the experimenter. C) Number of instances in which dyads engaged in verbal planning immediately before drawing together. D) Histograms of the number of utterances spoken by a dyad immediately before drawing together (verbal planning). Intergen = intergenerational group, Same gen = same generation group.

### External ratings

The external ratings (n = 12463 per rating scale, spread across 726 co-drawing drawings). The distributions of each rating scale are presented in Figure 11. These external ratings of drawings have been combined to make a measure of visible collaboration (summed use of space and coherence of motifs) and reported elsewhere ^59^.

**Figure 11.**
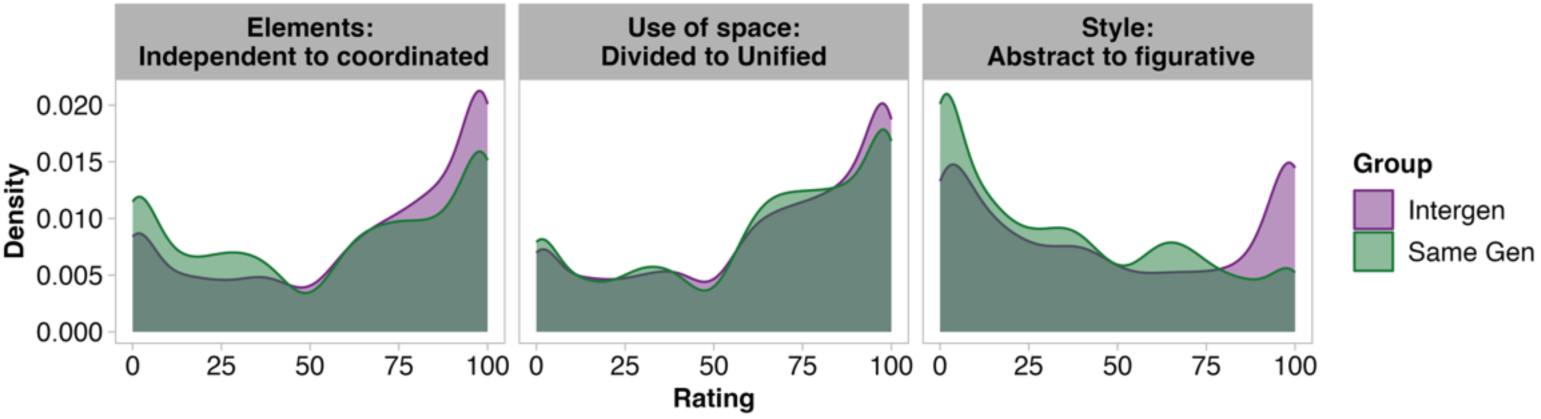
Ratings of drawing provided by external raters (unfamiliar with experiment) on the harmony between the elements in the drawings, the use of space in the drawing, and the artistic style of the drawing.

## 5 Usage Notes

Some technical difficulties were experienced during this study, resulting in a relatively small amount of missing data and some fNIRS recordings with unusual characteristics. The types of missing data and reasons that data are missing are detailed in Table 2. The fNIRS recordings with unusual characteristics, which require additional attention (or may be excluded from analyses), are listed in Table 3. As noted in the section on the technical validation of fNIRS data, there was a period during March 2023, when some right TPJ optodes were defective. This was identified based on SCIs of 1 with no cardiac oscillations. The issue was raised with Cortivision on March 4^th^, 2024, and Cortivision provided new optode bundles, which we began using on March 11^th^. We recommend excluding channels with no cardiac oscilations, as well as with SCI of 1, as we have done in our analyses of the fNIRS signals ^47^.

**Table 2.** A complete list of missing data. The list is organised by data type, with the reason the data is missing, as well as the specific participant, session, and dyad impacted.

| Data Type | Reason | Participant ID & Session | Dyad ID |
| --- | --- | --- | --- |
| All data types | Participant consumed alcohol prior to session | sub-243_ses-05 | 2026 (ses-05) |
| Majority of data | Participant unable to attend final session<br>(All data missing, except 6<sup>th</sup> collaborative activity and exit interview, which were collected at 5<sup>th</sup> session) | sub-139_ses-6<br>sub-140_ses-6<br>sub-317_ses-6<br>sub-319_ses-6 | 2017 (ses-06)<br>2019 (ses-06) |
| fNIRS | Recordings lost in transition between recording laptops | sub-101_ses-01<br>sub-102_ses-01<br>sub-103_ses-01<br>sub-124_ses-01<br>sub-201_ses-01<br>sub-202_ses-01<br>sub-211_ses-01<br>sub-301_ses-01 | 1001 (ses-01)<br>1002 (ses-01)<br>2001 (ses-01)<br>2006 (ses-01)<br>2015 (ses-01) |
| Motion capture | Video camera recording failure meaning no coordinates could be extracted from videos.<br>* indicates that no video recording was available and that no motion capture coordinates are included in the dataset<br>+ indicates that the video recording function for part of the session and that some motion capture coordinates are included in the dataset | sub-104_ses-05*<br>sub-105_ses-04*<br>sub-107_ses-02*<br>sub-203_ses-04*<br>sub-207_ses-02*<br>sub-212_ses-01*<br>sub-302_ses-05*<br>sub-308_ses-01*<br>sub-107_ses-03+<br>sub-109_ses-01+<br>sub-121_ses-03+<br>sub-123_ses-01+<br>sub-207_ses-03+<br>sub-209_ses-01+<br>sub-220_ses-03+<br>sub-323_ses-01+ | 1003 (ses-04)*<br>1005 (ses-02)*<br>2002 (ses-05)*<br>2008 (ses-01)*<br>1005 (ses-03)+<br>1007 (ses-01)+<br>1015 (ses-03)+<br>2023 (ses-01)+ |

**Table 3.** A list of fNIRS recordings that are included in the dataset but require special attention. The list details the reason the data requires special attention, as well as the specific participant, session, and dyad impacted.

| Reason | Participant ID<br>& Session | Dyad ID & Session |
| --- | --- | --- |
| Recordings used initial montage unoptimised TPJ coverage. Montage was updated as shown in Figure 4 for all other recordings. | sub-302_ses-01<br>sub-303_ses-01 | 2001 (ses-01)<br>2003 (ses-01) |
| Triggers missing. The research institutions network safety protocols occasionally interrupted triggers. | sub-137_ses-01<br>sub-140_ses-05<br>sub-206_ses-05<br>sub-236_ses-06<br>sub-304_ses-01<br>sub-304_ses-05<br>sub-319_ses-05 | 1029 (ses-01)<br>2004 (ses-01, ses-05)<br>2019 (ses-05) |
| Multi-part recording. The fNIRS signal was not recorded due Bluetooth signal dropped out or unanticipated low battery in the fNIRS device. | sub-123_ses-01<br>sub-216_ses-02<br>sub-221_ses-01<br>sub-227_ses-05<br>sub-231_ses-05<br>sub-310_ses-02 | 1016 (ses-01)<br>1020 (ses-05)<br>1024 (ses-05)<br>2010 (ses-02) |
| Signal disturbances caused by another wearable device, distributed by another research group at a local research institution. | sub-313_ses-02 | 2013 (ses-02) |

## Code Availability

The code for generating the validation values and figures can be found on the Open Science Framework (https://osf.io/g6z45).

## Acknowledgements

We would like to emphasise how thankful we are to the research and student assistants who helped collect and prepare the InterGenSynchrony Dataset. We thank Tessa Portier and Medea Häuselmann for their assistance with data collection. We thank Luca Naudszus for contributing to pose estimation. We thank Linda Fanconi for communicating with prospective participants and scheduling sessions. We thank Alistair Gadola for coding participant behaviour from videos. We thank Fanny Mougel for editing the digitised drawings. We thank Greta Bramow for creating the web-based experiment to collect external ratings and correcting text-based qualitative data. We thank Alistair Gadola and Filiz Kanele for correcting interview transcripts. We thank Ahmed Eldably for facilitating the transformation of the various data types into a BIDS compatible format. We thank Chris Markiewicz and the OpenNeuro Team for providing the platform and technical support needed to publish this dataset. We thank Leia Jekel for proofreading the manuscript.

## Author Contributions

RM: Conceptualization, equipment donation, data curation, formal analysis, investigation, visualization, writing – original draft. ESC: Conceptualization, funding acquisition, project administration, writing – review & editing.

## Competing Interests

The authors declare no potential competing interests.

## Funding

Two Cortivision Photon Cap devices were provided to RM through the Cortivision Pathfinder Program (grant number CPP-2023/09/01). Oil pastels were provided to RM by Caran D’Ache (agreement number 712517). RM and ESC were supported by the Social Brain Sciences Lab at ETH Zurich.

## Notes

https://openneuro.org/datasets/ds008192

### Competing Interest Statement

The authors have declared no competing interest.

### Summary of Updates

"hypercanning" in title corrected to "hyperscanning"

## References

1. Baumeister, R. F. & Leary, M. R. The need to belong: Desire for interpersonal attachments as a fundamental human motivation. in Interpersonal Development (eds Laursen, B. & Žukauskienė, R.) (Routledge, 2017). doi:10.4324/9781351153683.

2. Gable, S. L. Satisfying and meaningful close relationships. in The Social Psychology of Living Well (eds Forgas, J. P. & Baumeister, R. F.) 239–256 (Routledge, New York, NY: Routledge, 2018. | Series: The Sydney Symposium of Social Psychology series, 2018). doi:10.4324/9781351189712-14.

3. Leigh-Hunt, N. et al. An overview of systematic reviews on the public health consequences of social isolation and loneliness. Public Health 152, 157–171 (2017).

4. Rico-Uribe, L. A. et al. Association of loneliness with all-cause mortality: A meta-analysis. PLoS ONE 13, e0190033 (2018).

5. The Lancet. Loneliness as a health issue. The Lancet 402, 79 (2023).

6. WHO. Social Isolation and Loneliness among Older People: Advocacy Brief. https://www.who.int/publications/i/item/9789240030749 (2021).

7. WHO. From Loneliness to Social Connection. https://www.who.int/publications/i/item/978240112360 (2025).

8. Apostolou, M. & Keramari, D. What prevents people from making friends: A taxonomy of reasons. Personality and Individual Differences 163, 110043 (2020).

9. White, J. et al. Social connection, loneliness, and solutions: Perceptions of older adults. Activities, Adaptation & Aging 49, 671–695 (2025).

10. United Nations. World Population Prospects 2024: Summary of Results. (2024).

11. Moffat, R. & Cross, E. S. InterGenSynchrony Dataset. OpenNeuro (2026).

12. Holt-Lunstad, J. & Perissinotto, C. Social isolation and loneliness as medical issues. N Engl J Med 388, 193–195 (2023).

13. Scarpetti, G. et al. A comparison of social prescribing approaches across twelve high-income countries. Health Policy 142, 104992 (2024).

14. WHO. What Is the Evidence on the Role of the Arts in Improving Health and Well-Being? A Scoping Review. https://iris.who.int/handle/10665/329834 (2019).

15. Wigfield, A., Turner, R., Alden, S., Green, M. & Karania, V. K. Developing a new conceptual framework of meaningful interaction for understanding social isolation and loneliness. Social Policy and Society 21, 172–193 (2022).

16. Anderson, S. et al. Translating knowledge: Promoting health through intergenerational community arts programming. Health Promotion Practice 18, 15–25 (2017).

17. Jenkins, L. K., Farrer, R. & Aujla, I. J. Understanding the impact of an intergenerational arts and health project: A study into the psychological well-being of participants, carers and artists. Public Health 194, 121–126 (2021).

18. Lokon, E., Li, Y. & Kunkel, S. Allophilia: Increasing college students’ “liking” of older adults with dementia through arts-based intergenerational experiences. Gerontology & Geriatrics Education 41, 494–507 (2020).

19. Rubin, S. E. et al. Challenging gerontophobia and ageism through a collaborative intergenerational art program. Journal of Intergenerational Relationships 13, 241–254 (2015).

20. Lee, K., Jarrott, S. E. & Juckett, L. A. Documented outcomes for older adults in intergenerational programming: A scoping review: research. Journal of Intergenerational Relationships 18, 113– 138 (2020).

21. Newman, S., Ward, C., Smith, T., Wilson, J. & McCrea, J. Intergenerational Programs: Past, Present and Future. (Routledge, New York, 2014).

22. Petersen, J. A meta-analytic review of the effects of intergenerational programs for youth and older adults. Educational Gerontology 49, 175–189 (2023).

23. Bartlett, S. P., Solomon, P. & Gellis, Z. Comparative effectiveness of intergenerational service-learning programs on student outcomes of knowledge, attitude, and ageism. Educational Gerontology 47, 559–573 (2021).

24. Gonzales, E., Morrow-Howell, N. & Gilbert, P. Changing medical students’ attitudes toward older adults. Gerontology & Geriatrics Education 31, 220–234 (2010).

25. Dikker, S., Brito, N. H. & Dumas, G. It takes a village: A multi-brain approach to studying multigenerational family communication. Developmental Cognitive Neuroscience 65, 101330 (2024).

26. Zhang, Q., Liu, Z., Qian, H., Hu, Y. & Gao, X. Interpersonal competition in elderly couples: A functional near-infrared spectroscopy hyperscanning study. Brain Sciences 13, 600 (2023).

27. Liang, Z.-W., Zhan, Z.-J., Liu, Y.-C., Xu, H.-Z. & Yu, J. Interpersonal neural synchronization underlies interactive concept learning in older adults. *npj Sci*. Learn. 10, 75 (2025).

28. Bevilacqua, D. et al. Brain-to-brain synchrony and learning outcomes vary by student–teacher dynamics: Evidence from a real-world classroom electroencephalography study. Journal of Cognitive Neuroscience 31, 401–411 (2019).

29. Dikker, S. et al. Brain-to-brain synchrony tracks real-world dynamic group interactions in the classroom. Current Biology 27, 1375–1380 (2017).

30. Ellingsen, D.-M. et al. Brain-to-brain mechanisms underlying pain empathy and social modulation of pain in the patient-clinician interaction. Proc. Natl. Acad. Sci. U.S.A. 120, e2212910120 (2023).

31. Grahl, A. et al. Brain and behavioral correlates of the patient-clinician relationship: A longitudinal fMRI hyper-scanning study of chronic pain patients. The Journal of Pain 22, 602 (2021).

32. Grahl, A. et al. The patient-clinician relationship, expectancy and prior experience can modulate fibromyalgia treatment outcomes: a longitudinal fmrihyperscan study. The Journal of Pain 24, 74–75 (2023).

33. Francis, M., Markus, A., Stuhr-Wulff, F. & Shamay-Tsoory, S. Shaping inter-brain plasticity: A feasibility study of enhancing inter-brain synchrony with dyadic neurofeedback. iScience 29, 114894 (2026).

34. Moffat, R., Casale, C. E. & Cross, E. S. Mobile fNIRS for exploring inter-brain synchrony across generations and time. Frontiers in Neuroergonomics 4, (2024).

35. Lakens, D. Sample Size Justification. Collabra: Psychology 8, 33267 (2022).

36. Paulus, C. Der Saarbrücker Persönlichkeitsfrageboden SPF (IRI) zur Messung von Empathie: Psychometrische Evaluation der deutschen Version des Interpersonal Reactivity Index. (2009) doi:10.23668/psycharchives.9249.

37. Davis, M. A multidimensional approach to individual differences in empathy. Journal of Personality and Social Psychology 10, 85 (1980).

38. De Jong Gierveld, J. & van Tilburg, T. A 6-item scale for overall, emotional, and social loneliness: Confirmatory tests on survey data. Res Aging 28, 582–598 (2006).

39. Pittinsky, T. L., Rosenthal, S. A. & Montoya, R. M. Measuring positive attitudes toward outgroups: Development and validation of the Allophilia Scale. in Moving beyond prejudice reduction: Pathways to positive intergroup relations. (eds Tropp, L. R. & Mallett, R. K.) 41–60 (American Psychological Association, Washington, 2011). doi:10.1037/12319-002.

40. Aron, A., Aron, E. N. & Smollan, D. Inclusion of other in the self scale and the structure of interpersonal closeness. Journal of Personality and Social Psychology 63, 17 (1992).

41. Peirce, J. W., Hirst, R. J. & MacAskill, M. R. Building Experiments in PsychoPy. (Sage, London, 2022).

42. Oostenveld, R. & Praamstra, P. The five percent electrode system for high-resolution EEG and ERP measurements. Clinical Neurophysiology 112, 713–719 (2001).

43. Zimeo Morais, G. A., Balardin, J. B. & Sato, J. R. FNIRS Optodes’ Location Decider (fOLD): A toolbox for probe arrangement guided by brain regions-of-interest. Scientific Reports 8, 1–11 (2018).

44. Oostenveld, R., Fries, P., Maris, E. & Schoffelen, J.-M. Fieldtrip: open source software for advanced analysis of MEG, EEG, and invasive electrophysiological data. Computational Intelligence and Neuroscience 2011, 1–9 (2011).

45. Cao, Z., Hidalgo, G., Simon, T., Wei, S.-E. & Sheikh, Y. Openpose: Realtime multi-person 2d pose estimation using part affinity fields. IEEE Trans. Pattern Anal. Mach. Intell. 43, 172–186 (2021).

46. BIDS structure for longitudinal dyadic data - Neuro Questions. Neurostars https://neurostars.org/t/bids-structure-for-longitudinal-dyadic-data/26173 (2023).

47. Moffat, R., Dumas, G. & Cross, E. S. Longitudinal hyperscanning indexes changes in social connection in intergenerational and same generation dyads. PLOS Biology (In Press) doi:10.1101/2025.10.14.682029.

48. De Felice, S., et al. Having a chat and then watching a movie: how social interaction synchronises our brains during co-watching. Oxford Open Neuroscience 3, kvae006 (2024).

49. Cui, X., Bryant, D. M. & Reiss, A. L. NIRS-based hyperscanning reveals increased interpersonal coherence in superior frontal cortex during cooperation. NeuroImage 59, 2430–2437 (2012).

50. Czeszumski, A. et al. Cooperative behavior evokes interbrain synchrony in the prefrontal and temporoparietal cortex: A systematic review and meta-analysis of fNIRS hyperscanning studies. eNeuro 9, ENEURO.0268-21.2022 (2022).

51. Dommer, L., Jäger, N., Scholkmann, F., Wolf, M. & Holper, L. Between-brain coherence during joint n-back task performance: A two-person functional near-infrared spectroscopy study. Behavioural Brain Research 234, 212–222 (2012).

52. Li, L. et al. Interpersonal neural synchronization during cooperative behavior of basketball players: a fNIRS-based hyperscanning study. Frontiers in Human Neuroscience 14, (2020).

53. Osaka, N. et al. How two brains make one synchronized mind in the inferior frontal cortex: fNIRS-based hyperscanning during cooperative singing. Frontiers in Psychology 6, (2015).

54. Sun, B. et al. Cooperation with partners of differing social experience: An fNIRS-based hyperscanning study. Brain and Cognition 154, 105803 (2021).

55. Xie, H. et al. Finding the neural correlates of collaboration using a three-person fMRI hyperscanning paradigm. Proc. Natl. Acad. Sci. U.S.A. 117, 23066–23072 (2020).

56. Moffat, R., Dumas, G. & Cross, E. S. Longitudinal intergenerational hyperscanning indexes changes in social connection. 2025.10.14.682029 Preprint at 10.1101/2025.10.14.682029 (2026).

57. Zhang, M. et al. Multi-view emotional expressions dataset using 2D pose estimation. Sci Data 10, 649 (2023).

58. Cronbach, L. J. Coefficient alpha and the internal structure of tests. Psychometrika 16, 297–334 (1951).

59. Moffat, R., Naudszus, L. A. & Cross, E. S. Cardiac synchrony during collaborative drawing: A longitudinal comparison of same generation and intergenerational dyads. Ann NY Acad Sci 1558, e70272 (2026).

60. Naudszus, L. A., Moffat, R. & Cross, E. S. Two-brain states characterise within-and between-brain connectivity during intergenerational collaborative drawing. Acta Psychologica (In Press) doi:10.64898/2026.01.29.702550.

